# Maturation-dependent changes in cortical and thalamic activity during slow waves of light sleep: insights from a combined EEG-fMRI study

**DOI:** 10.1101/2023.06.07.542852

**Authors:** Damiana Bergamo, Giacomo Handjaras, Flavia Petruso, Francesca Talami, Emiliano Ricciardi, Francesca Benuzzi, Anna Elisabetta Vaudano, Stefano Meletti, Giulio Bernardi, Monica Betta

## Abstract

**INTRODUCTION:** Studies using scalp EEG have shown that slow waves (0.5-4 Hz), the most prominent hallmark of NREM sleep, undergo relevant changes from childhood to adulthood, mirroring brain structural modifications and the acquisition of cognitive skills. Here we used simultaneous EEG-fMRI to investigate the cortical and subcortical correlates of slow waves in school-age children and determine their relative developmental changes.

**METHODS:** We analyzed data from 14 school-age children with self-limited focal epilepsy of childhood who fell asleep during EEG-fMRI recordings. Brain regions associated with slow-wave occurrence were identified using a voxel-wise regression that also modeled interictal epileptic discharges and sleep spindles. At the group level, a mixed-effects linear model was used. The results were qualitatively compared with those obtained from 2 adolescents with epilepsy and 17 healthy adults.

**RESULTS:** Slow waves were associated with hemodynamic-signal decreases in bilateral somatomotor areas. Such changes extended more posteriorly relative to those in adults. Moreover, the involvement of areas belonging to the default mode network changes as a function of age. No significant hemodynamic responses were observed in subcortical structures. However, we identified a significant correlation between age and thalamic hemodynamic changes.

**CONCLUSIONS:** Present findings indicate that the somatomotor cortex may have a key role in slow-wave expression throughout the lifespan. At the same time, they are consistent with a posterior-to-anterior shift in slow-wave distribution mirroring brain maturational changes. Finally, our results suggest that slow-wave changes may not reflect only neocortical modifications but also the maturation of subcortical structures, including the thalamus.

**Statement of significance:** Low spatial resolution of conventional EEG has widely limited the possibility of accurately determining in humans the role of different brain structures in the expression of sleep slow waves throughout development. Here, for the first time, we took advantage of a simultaneous EEG-fMRI approach to accurately describe the cortical and subcortical hemodynamic correlates of sleep slow waves in a sample of 14 school-aged children. In order to elucidate age-dependent changes, we qualitatively compared present findings with those previously obtained in a group of adults. Overall our results have important implications for the understanding of how cortico-cortical and subcortico-cortical interactions shape sleep slow waves across development.

## Introduction

The sleep slow wave (<4.5 Hz) represents the main electroencephalographic (EEG) hallmark of human non-Rapid-Eye-Movement (NREM) sleep. Sleep slow waves arise from the synchronized oscillation of large neuronal populations between a hyperpolarized state of ‘silence’ (*off-period*) and a depolarized, ‘active’ state (*on-period*)^1,2^. Even though slow waves involve and are generated at the cortical level, a growing body of evidence indicates that subcortical structures, including several brainstem nuclei and the thalamus, have a crucial role in regulating their physiological expression ^3–5.^ Indeed, recent studies based on simultaneous thalamic and cortical recordings in humans, showed that changes in thalamic activity accompany the occurrence of cortical slow waves and may have an active role in determining their coordination with other sleep rhythms, such as sleep spindles (10-16 Hz)^6,7^. Consistent with this, studies combining EEG and functional magnetic resonance imaging (fMRI) described an association between cortical EEG slow waves and hemodynamic changes in subcortical structures, including the thalamus and distinct portions of the brainstem^8–10^.

While sleep was classically regarded as a global state of the whole brain, it is now well established that most slow waves occur and are regulated locally, as a function of prior experience and learning ^11–15^. Thus, brain areas that are activated and involved in learning during wakefulness typically show increased slow-wave activity (SWA; power in the delta range, 1-4.5 Hz) during subsequent sleep. Such changes may reflect an increased local ‘sleep need’, which has been hypothesized to depend on the necessity to normalize experience-dependent increases in synaptic strength^16,17^ and/or counteract the accumulation of metabolic wastes resulting from neuronal activity^18–20^. Moreover, slow waves travel at cortical level following cortico-cortical connections and are thus regarded as a valuable read-out of long-range functional and anatomical connectivity^21–23^.

Interestingly, the morphological and topographical features of slow waves also appear to change as a function of brain development, from childhood to adulthood^24^. Indeed, a seminal high density EEG (hd-EEG) study by Kurth and colleagues25 demonstrated that the scalp location of maximal SWA moves from centro-posterior regions in early childhood to frontal areas in late adolescence. This posterior-to-anterior shift has been suggested to mirror anatomo-functional changes related to brain maturation^26–28^-including synaptic pruning and redistribution^29,30^-, and the acquisition of specific cognitive and behavioral skills^31,32^. Moreover, slow-wave spread and traveling distance increase from childhood to adulthood, likely reflecting the maturation of (long-range) anatomical wiring patterns^33,34^. Of note, though, the potential contribution of subcortical structures - and the thalamus, in particular - to maturational changes in the expression of human sleep slow waves has never been directly investigated.

Using simultaneous EEG-fMRI, we recently identified brain regions showing hemodynamic signal changes during EEG slow waves in adults^8^. Specifically, we demonstrated that cortical slow waves of light sleep are associated both with negative hemodynamic signal changes in the somatomotor, visual, and insular cortices and with positive signal changes in subcortical structures including the thalamus, brainstem, and cerebellum. Here we used the same approach to investigate the cortical and subcortical hemodynamic correlates of sleep slow waves and their relative developmental modifications in a sample of school-age children with a diagnosis of self-limited focal epilepsy of childhood who fell asleep during an afternoon resting-state scan session^35^.

## Methods

### Participants

The present study was based on a previously collected simultaneous EEG-fMRI resting-state dataset obtained from school-age children and adolescents with a diagnosis of self-limited focal epilepsy of childhood, including in particular self-limited epilepsy with centrotemporal spikes (SeLECTS) and childhood occipital visual epilepsy (COVE)^35–37^. Self-limited focal epilepsies refer to a group of epileptic conditions characterized by seizures originating from a specific region of the brain and having, in most of the cases, a favorable prognosis for spontaneous resolution over time without the need for long-term medication or intervention^35^.

Recordings were performed with the aim of investigating the fMRI correlates of epileptic activity during wakefulness^36,37^. Nevertheless, some study participants fell asleep during one or more of the undertaken scan sessions. Thus, for the present study, a total of 27 subjects with SeLECTS and 9 subjects with COVE were initially selected as they showed signs of potential sleep based on visual inspection of their EEG traces. Then, the EEG recordings were preprocessed as described below and re-assessed systematically for the presence of sleep and artifactual activity. This procedure led to the exclusion of 20 subjects (no or insufficient sleep, N=9; presence of strong residual artifactual activity, N=11), of which 8 with COVE and 12 with SeLECTS.

Analyses described in the present study were based on a final sample of 14 school-age children (2 females; mean age ± SD = 8.4±1.6, range 6-11 yrs) with a diagnosis of SeLECTS. As detailed below, data obtained from two older adolescents (1 female; age 15 yrs and 17 yrs), respectively diagnosed with SeLECTS and COVE, were used for qualitative comparisons in analyses aimed at assessing thalamic activity associated with the occurrence of sleep slow waves. Six out of fourteen children and both adolescents were under antiepileptic medication at the time of the EEG-fMRI recording (Table 1). As per definition of Self-limited childhood epilepsy, all patients had no diagnosis of other neurologic disorders besides epilepsy and no pathological abnormalities were identified on conventional MRI. The study was approved by the Local Ethic Committee of the Azienda Ospedaliera Universitaria di Modena (80/10). Written informed consent was obtained from parents and assent from patients. Hemodynamic cortical and thalamic responses observed in children with SeLECTS were compared qualitatively with those obtained from a sample of 17 healthy adults (11 females; mean age ± SD = 28.8 ± 2.3, range 25–35 yrs) acquired during an afternoon nap using the same MRI scanner and EEG system and described in our previous work^8^.

**Table 1.** Demographic and sleep-related variables. All analyses focused on 14 children with age comprised between 6 and 11 yrs (left column). For qualitative comparison, we also reported results obtained from two adolescents (right column). SeLECTS = self limited epilepsy with Centro Temporal Spikes; COVE = Childhood Occipital Visual Epilepsy; ASM = antiseizure medicines; VPA = Valproic Acid; LEV = Levetiracetam; CLB = Clobazam; OXC = Oxcarbazepine; CBZ = Carbamazepine; LTG = Lamotrigine; LCS = Lacosamide; ZNS = Zonisamide.

| Age groups (n=16) | Children (6-11 y) |  | Adolescents (15-17 y) |  |
| --- | --- | --- | --- | --- |
| Group Variables | N / mean | STD | N / mean | STD |
| **N** | 14 | X | 2 | X |
| **M / F** | 12/2 | X | 1/1 | X |
| **Age** | 8.4 | ± 1.3 | 16 | ± 1.4 |
| **Handedness** | 13 right | X | 2 right | X |
| **Diagnosis** | 14 self-limited focal epilepsy (SeLECTS) | X | 2 self-limited focal epilepsy (1 SeLECTS, 1 COVE) | X |
| **ASM** | VPA (2), LEV (2), CLB (1), OXC (1), CBZ (2) | X | LTG (1), LCS (1), ZNS (1), CLB (1), VPA (1) | X |
| Sleep Variables | min (%) / N | STD | min (%) / N | STD |
| **Wake** | 8.50 (4.44%) | ± 1.02 | 2 (10%) | ± 0.71 |
| **NREM** | 183 (95.56%) | ± 5.67 | 18 (90%) | ± 4.06 |
| **N1 (%)** | 82.50 (43.08%) | ± 6.87 | 11.5 (57.5%) | ± 5.31 |
| **N2 (%)** | 66 (34.46%) | ± 4.69 | 6.5 (32.5%) | ± 4.60 |
| **N3 (%)** | 34.50 (18.02 %) | ± 5.08 | 0 (0%) | ± 0 |
| **Slow waves (N per participant)** | 130.64 (11.91 SW/min) | ± 91.01 | 52.50 (5.67 SW/min) | ± 27.57 |
| **IEDs (N per participant)** | 729.14 (45.38 IEDs/min) | ± 801.50 | 617 (74.69 IEDs/min) | ± 517.60 |
| **Spindles (N per participant)** | 17.92 (1.54 spd/min) | ± 18.79 | 4.5 (0.69 spd/min) | ± 6.36 |

### EEG-fMRI Protocol

Simultaneous EEG-fMRI recordings were performed in the early afternoon. No sleep deprivation or sedation protocols were used. To increase the comfort of the subjects inside the scanner and minimize head movements, secure EEG leads and foam pads were used. Subjects were asked to remain still during the scanning with their eyes closed. Further details regarding the clinical assessment of subjects have been described in previous work^36^.

Functional and anatomical MRI data were acquired using a 3T Philips Intera system. T2*-weighted gradient-echo echo-planar sequences were used to acquire data from 30 contiguous axial slices over 8-minute (TR = 2,000 ms; in-plane matrix = 80 × 80; voxel size = 3 × 3 × 4 mm; 240 volumes) or 10-minute runs (TR = 3,000 ms; in-plane matrix = 64 × 64; voxel size = 4 × 4 × 4 mm; 200 volumes).

A high-resolution T1-weighted anatomical image was also obtained in all subjects (TR = 9.9ms; TE = 4.6ms; 170 sagittal slices; in-plane matrix = 256 × 256; voxel size = 1 × 1 × 1 mm).

Scalp EEG was recorded using a 32-channel MRI-compatible EEG recording system (Micromed, Mogliano Veneto, Italy). EEG channels were placed according to conventional 10-20 locations, and sampled at 1024 Hz. Data was transmitted via an optic fiber cable from the amplifier to a computer located outside the scanner room. EEG amplifiers had a resolution of 22 bits with a range of ± 25.6 mV.

### EEG Preprocessing

All EEG recordings were preprocessed in MATLAB (The MathWorks, Inc., version 2017a) using EEGLAB^38^ and following the same procedure described in previous work^8^. Specifically, we used the FMRIB plugin^39,40^ to remove fMRI-related artifacts from EEG data. The algorithm removes MR gradient artifacts based on average artifact subtraction (AAS), subtracts the principal components (PC) of artifact residuals (Optimal Basis Sets method; OBS), and applies an adaptive noise cancellation (ANC). The signal was then down-sampled to 256 Hz. The removal of ballistocardiography artifacts followed an automated detection of QRS complexes in the ECG channel and was based on the subtraction of PCs of artifact residuals (OBS method). All automated QRS detections were visually inspected and wrong or missing markers were manually corrected using custom-made MATLAB functions and according to criteria described in previous work^41^. Finally, EEG recordings were band-pass filtered between 0.4 and 35 Hz prior to the application of an Independent Component Analysis (ICA)^38^, aimed at reducing any residual physiological artifacts. Bad channels were visually identified, rejected, and interpolated using spherical splines from the activity of the nearest sensors. The EEG signal was re-referenced to the average of channels T5 and T6 (representing the closest electrodes to the ASSM-recommended mastoid references) and sleep scoring was performed over 30-s epochs according to standard criteria^42^. We retained for the present analyses only recordings that included a minimum amount of NREM (N1/N2/N3) sleep corresponding to 50% of the run duration. Body movements and isolated artifacts were manually marked upon visual inspection of the EEG signals and were excluded from further analyses. Interictal epileptiform discharges (IEDs) were manually marked by two expert neurophysiologists^36^.

### Slow-wave detection

Slow waves were detected using a validated algorithm based on the identification of signal zero-crossings in an artificial EEG channel representing the negative signal envelope computed across all available electrodes^43^. Differently from commonly applied channel-by-channel detection approaches, this method allows for the identification of both local and widespread slow waves and defines a unique time reference (across electrodes) for each negative wave^44^. In particular, the negative-going envelope of the 0.5-18 Hz filtered signal was calculated for each time-point by selecting the four most negative samples (across electrodes), discarding the single most negative sample, and taking the average of remaining values. This approach was used to minimize the potential impact of any residual large-amplitude artifactual activity in isolated electrodes. The resulting signal was baseline corrected (zero-mean centered) before applying a negative-half-wave detection based on the identification of consecutive signal zero-crossings^43,45^. Only negative half-waves detected during NREM sleep epochs (N1/N2/N3) and with a duration comprised between 0.25 s and 1.0 s (full-wave period 0.5–2.0 s, corresponding to a 0.5–2.0 Hz frequency range) were selected for subsequent analyses. No amplitude thresholds were applied ^8,23,46,47^.

### Spindle detection

Since spindles often occur in association with slow waves, we assessed their possible impact on hemodynamic signal changes during slow waves. As previously described, spindles were automatically detected in the signal of channel Cz using a validated algorithm^48,49^. Specifically, a wavelet-based filter (10–16 Hz) was applied to the EEG time-series using a b-spline wavelet^50^, and the time course of power of the resulting signal was measured by squaring the values and smoothing the time-series using a sliding window of 100 ms. Then, potential spindles were defined as points in which power values passed a high-threshold corresponding to the median plus 4 times the median absolute deviation (MAD) of signal power. The actual start and end points of identified events were then measured using the crossing times at a second, low-threshold corresponding to the median power plus 2 MADs. The thresholds were re-computed for each 30-s epoch. Only events with duration between 0.3 and 3 s and detected during N2/N3 sleep or transitional N1 epochs (i.e., N1 epochs adjacent to N2/N3 epochs) were retained for further analyses^8^. Finally, a power-ratio threshold was applied in order to ensure some specificity of the transient power increases within the spindle range. Specifically, for each potential spindle, we computed the ratio of the mean power in the spindle range (10-16 Hz) over the mean power in the neighboring ranges (8–10 Hz and 16–18 Hz) and we eventually retained the detected event if the obtained value was greater than 3.

### (f)MRI Data Preprocessing

Preprocessing of MRI and fMRI signals was performed using AFNI (version 19.1.16)^51^, FMRIB Software Library (FSL, version 5.0.9)^52^, Statistical Parametric Mapping (SPM-12, version 12.6906, https://www.fil.ion.ucl.ac.uk/spm/)^53^ and Advanced Normalization Tools (ANTs, http://stnava.github.io/ANTs/, version 2.1.0)^54^.

First, a bias-field correction of the T1w images was applied using SPM. The T1w data was then skull-stripped with ANTS, aligned to match the EPI (*align_epi_anat*.*py*), and non-linearly warped (*3dQwarp*) using as base a pediatric atlas from the NIH research center (https://nist.mni.mcgill.ca/pediatric-atlases-4-5-18-5y/)^55^.

Functional images were corrected for signal outliers using *3dDespike* and temporally aligned with *3dTshift*. Data from different runs were registered to the mean of the run for motion correction using *3dvolreg*. In addition, we used *fsl_motion_outliers* to detect timepoints corrupted by large movements. We then spatially smoothed the data (*3dBlurToFWHM*) with a 6 mm full-width-at-half-maximum (FWHM) Gaussian kernel and computed for each voxel the percent of Blood-Oxygen-Level-Dependent (BOLD) signal change with respect to the signal mean. Data were non-linearly transformed (*3dNwarpApply*) into the NIH pediatric atlas coordinate system and resampled to a 3-mm iso-voxel resolution. Finally, we regressed out from the BOLD signal of each voxel (*3dDeconvolve*) head-motion parameters, movement spike regressors (framewise displacement above 0.3), the mean signal of cerebrospinal fluid (CSF)^56^, and the temporal autoregression (ARMA-1, *3dREMLfit*), which typically reflects artifacts of physiological origin^57^.

### EEG-based regression analysis of fMRI time-series

At a within-subject level, the regions associated with slow-wave occurrence were identified through a voxel-wise regression analysis of BOLD time-series. Considering the possible temporal association between sleep spindles and slow waves, as well as the potential impact of IEDs on the BOLD signal, both spindles and IEDs were modeled in the regression. We built the slow-wave regressor considering each slow wave as a square wave with onset-time corresponding to the first zero-crossing of the negative half-wave, duration equal to the duration of the descending phase of the slow wave (from the first zero-crossing to the maximum negative peak) and amplitude equal to the absolute value of the maximum negative peak^8^. A similar procedure was used for the construction of the spindle regressor. In this case, each spindle was modeled as a binary square wave with onset time and duration derived from the automated detection algorithm. The obtained regressors were then convoluted with a standard gamma hemodynamic response function and down-sampled to the BOLD time-series sampling rate. Finally, a regressor describing epileptic activity was obtained by computing for each TR the IED density (number of detected IEDs divided by the length of the TR) and performing a convolution with a standard gamma response function.

The regressors and each BOLD time-series were forced to zero and the baseline value, respectively, in correspondence to artifactual and wakefulness data segments. At the single-subject level, beta-values of each cortical voxel were converted into z-scores calculated with respect to a null distribution of beta values obtained after randomly shuffling the timing of all the EEG events of interest (e.g., slow waves, spindles, IED density; intra-voxel and intra-subject permutations; N = 1000). The total number of slow waves and spindles and the IED density values were kept constant across original and shuffled regressors. Moreover, the repositioning of EEG events within artifactual and wakefulness data segments was prevented.

For each voxel, a mixed-effect model was used for assessing statistical significance at the group-level. In the model, z-scores obtained from each fMRI run were taken as the response variable while subjects were considered as the grouping variable. We then applied a cluster-based correction procedure to identify significant clusters and to account for multiple comparisons across voxels (cluster-forming threshold = p<0.01, cluster-wise significance threshold p <0.05; 1000 permutations). Figure 1 summarizes the main steps applied for the voxel-wise regression analysis.

**Figure 1.**
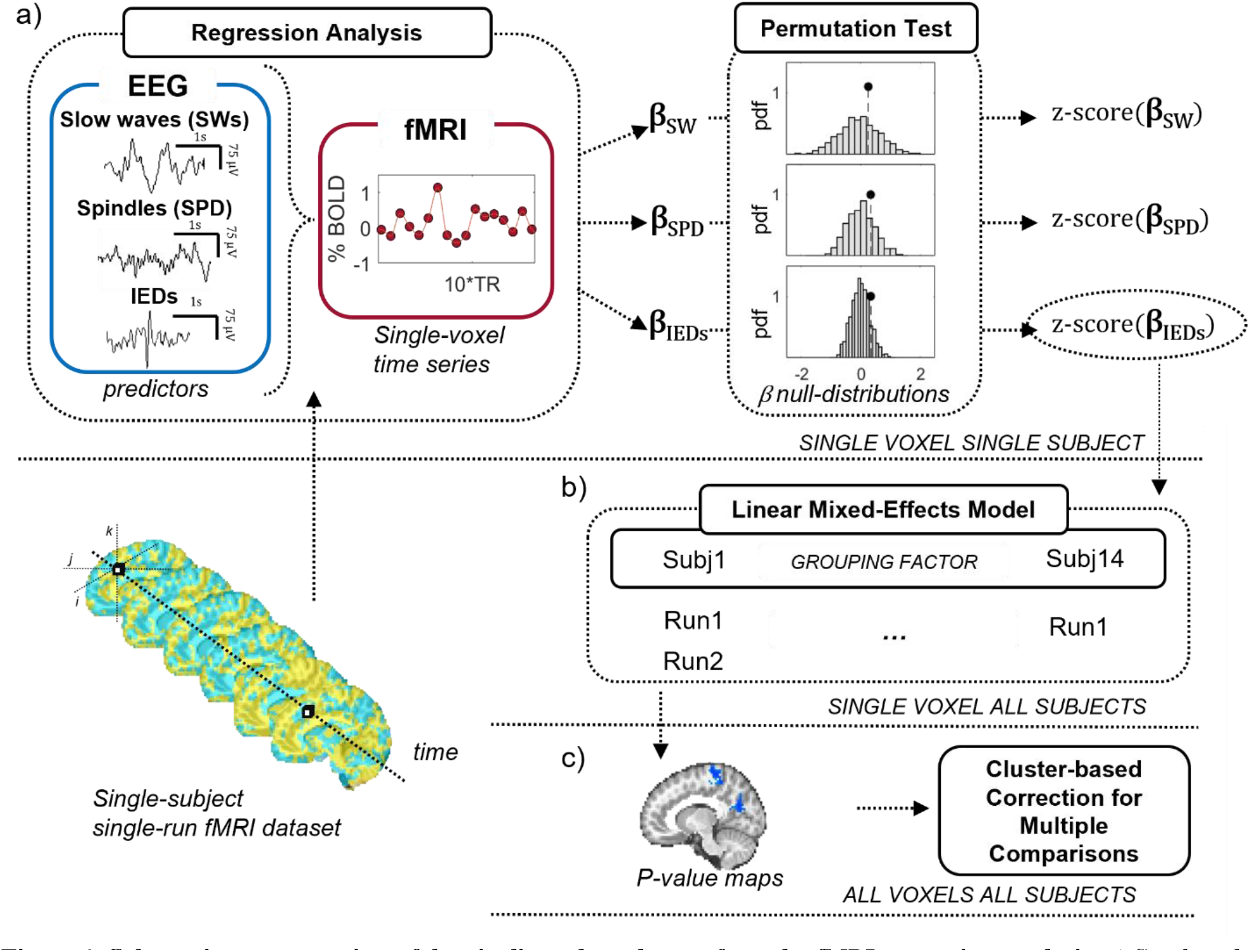
Schematic representation of the pipeline adopted to perform the fMRI regression analysis. a) Single-subject, single-voxel regression of BOLD time-series modeling as predictors slow-waves, spindles and IEDs. Obtained beta values associated with each regressor were z-scored considering a correspondent null distribution generated by shuffling the timing of individual events (n=1000 permutations). b) Group-level analysis based on a voxel-wise linear mixed effects model considering subjects as a grouping factor. c) Results of the group-level analysis were corrected using a cluster-based correction for multiple comparisons (cluster-forming threshold=p<0.01, cluster-wise significance threshold p<0.05).

### Comparison of cortical slow-wave-related hemodynamic changes in children and adults

The distribution of scalp SWA is known to shift from posterior to anterior electrodes from childhood to adulthood^25^. To investigate the hemodynamic correlates of this developmental change, cortical activity maps obtained in children were qualitatively compared with those previously obtained using a similar experimental protocol and analysis in a group of 17 healthy adults^8^. Since the present and the previous study relied on different degrees of freedom and adopted different correction methods to account for multiple statistical testing, the comparison was performed using the uncorrected statistical maps obtained from the voxel-wise regression analysis. Specifically, we thresholded group-level cortical maps of children and adults at p<0.01. In order to compare the two maps, the one with the largest number of voxels was further refined considering only the N most significant voxels, where N is the number of voxels included in the other map. Namely, the cortical involvement related to slow waves was kept constant across the two groups while assessing the spatial distribution of the most significant voxels. The percentage of cortical voxels belonging to each consecutive coronal slice of the resulting dataset, in the posterior to anterior direction, was computed for both adult and children maps, as well as for their overlap.

### Effect of age on cortical regional involvement in sleep slow waves

Slow waves behave as traveling waves, propagating at the cortical level through anatomically connected pathways^21,22^. Consistent with this, we recently showed in adults that hemodynamic changes associated with slow waves can extend, although with slightly different relative delays, to a broad portion of the entire cortex^8^. In light of known developmental changes in slow-wave cortical synchronization and propagation, here we investigated the possible effect of age on coordinated hemodynamic changes within different cortical networks during slow waves. Specifically, for each subject, we computed the average BOLD profiles time-locked to slow-wave onset within each of the 200 cortical regions of interest (ROIs) defined in the Schaefer functional atlas^58^. Average BOLD profiles were computed in a temporal interval between 0 and 14 seconds after slow wave onset. Of note, the IED density contribution estimated with the regression was subtracted from the BOLD signal of each voxel to minimize the potential effect of epileptic activity on signal changes. Then, the ROIs were assigned to one of seven functional networks^59^: visual, somatomotor, dorsal attention, ventral attention, limbic, frontoparietal, and default mode (DMN) networks. The overall degree of similarity among the signal temporal profiles of ROIs belonging to each network was assessed averaging all positive values within the pairwise correlation coefficients among each pair of ROIs. Finally, the correlation between within-network signal similarity and age was computed for each of the seven canonical networks. The significance of the results was tested using a non-parametric permutation test, in which the same correlation was computed with an identical procedure after randomly shuffling the timing of individual slow waves for each subject (N = 1000).

### Correlation between thalamic slow-wave-related hemodynamic changes and age

Previous work conducted on animal models^5,60^ and studies on human sleep based on intracranial recordings^7^ and EEG-fMRI^8^ are consistent with a role of the thalamus in slow-wave generation and/or regulation. We thus investigated whether the thalamic contribution to slow waves might show maturation-dependent modifications. For each subject, we first regressed out from the BOLD signal of each voxel the estimated contribution of the IED-based regressor to remove the effect of epileptic activity. Then, thalamic voxels were assigned to seven ROIs based on their preferential functional connectivity to one of the canonical cortical networks^59^. This functional parcellation of the thalamus was based on resting-state fMRI data collected in a group of 28 adults (range 19-43 yrs, for details see ^8^) and was also adopted in our previous work investigating the hemodynamic correlates of sleep slow waves in adults. For each ROI, we averaged across voxels the BOLD-signal changes time-locked to the onset of slow waves in a temporal interval between-4 up to 16 seconds after wave onset. Then, we identified the maximum value (positive peak) occurring between 3 and 5 seconds after slow-wave onset. This particular time interval was chosen as it corresponds to the timespan in which the peak of the hemodynamic response can be expected to occur based on the standard hemodynamic response function. Finally, we performed a correlation between the BOLD-signal maximum amplitude and age for each thalamic ROI. The statistical significance of the results was assessed using a non-parametric permutation test. Specifically, the same correlation was computed after randomly shuffling the timing of individual slow waves for each subject (N = 1000). This procedure generated a null distribution of correlation values which were used to calculate the quantile of the true correlation coefficient.

## Results

### Sample and sleep macrostructure

The final sample (Table 1) consisted of 14 school-age children (6-11 yrs; one 8-minute run for 2 participants, one 10-minute run for 5 participants, two 8-minute runs for 1 participant, one 8-minute run and one 10-minute run for 5 participants and two 10-minute runs for 1 participant) and two older adolescents (15-17 yrs; one 10 min run for each participant). The sleep of participants included, on average, 13.07 minutes (± 4.61, range 7-20 min) of NREM (N1= 5.89 ± 6.87 min; N2= 4.71 ± 4.69 min; N3= 2.46 ± 5.08 min). The mean power spectral density from all the analyzed EEG runs is shown in Figure 2a, together with a representative 30-second EEG trace recorded during N2 sleep (Figure 2b). A total of 1829 slow waves (130.64 ± 91.01 per participant), 10208 IEDs (729.14 ± 801.50 per participant), and 251 sleep spindles (17.92 ± 18.79 per participant) were detected and included in the analyses.

**Figure 2.**
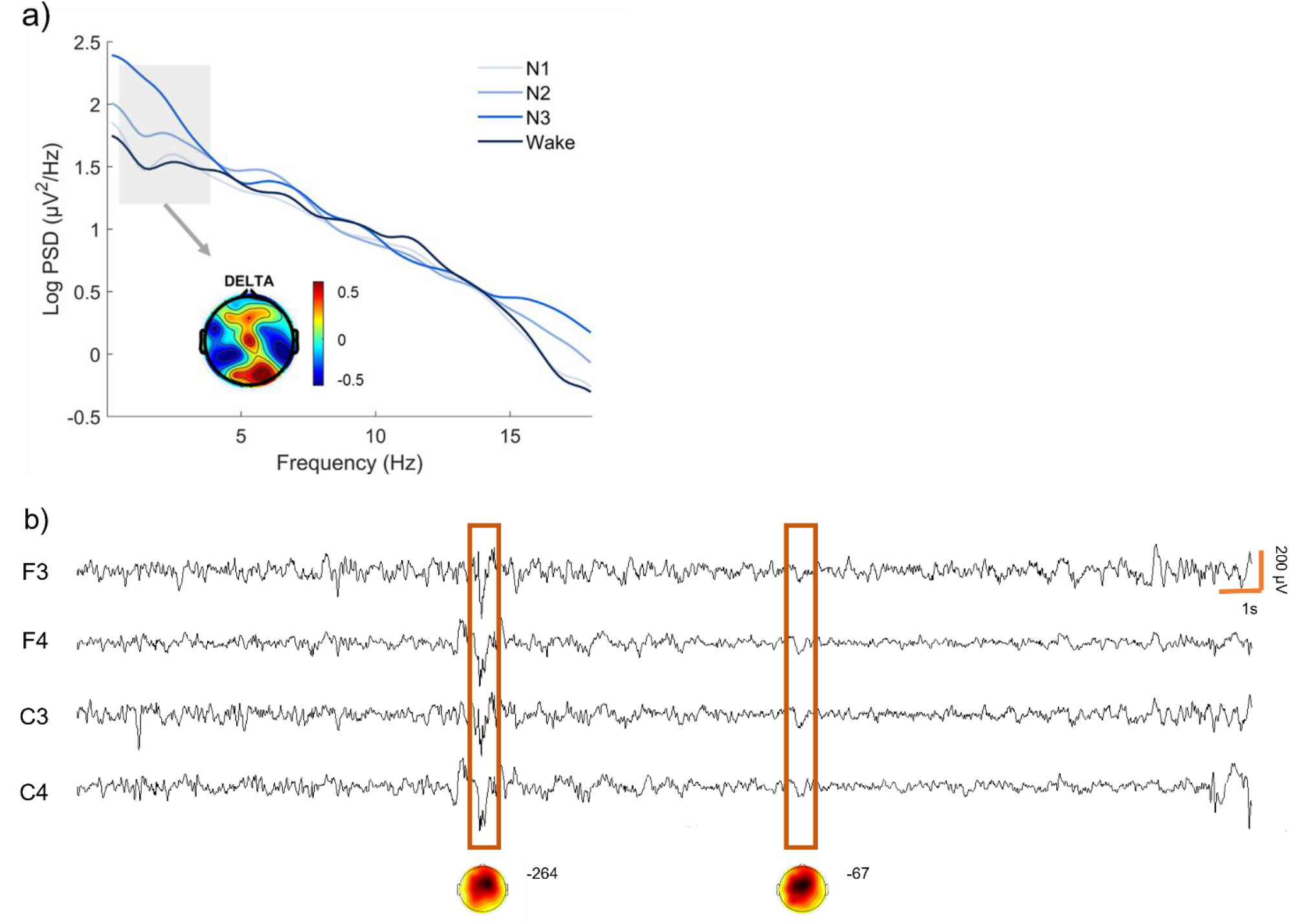
Analysis of the EEG recording. a) Mean power spectral density obtained across all school-age participants and all analyzed runs, divided by sleep stage (Wake, N1, N2, N3). Spectral density values were calculated in two frontal (F3, F4) and two central (C3, C4) electrodes and then averaged. The topographic distribution of delta power is also shown. b) Representative EEG traces from an N2 epoch. Detected slow waves are highlighted with orange boxes. The corresponding topographical distribution is shown below the boxes together with the maximum amplitude of the wave negative peak (in µV).

### Hemodynamic signal changes associated with slow waves

In order to explore the cortical and subcortical hemodynamic correlates of sleep slow waves in school-age children, we performed a voxel-wise multiple regression analysis of the hemodynamic BOLD signal by modeling slow waves, IEDs, and sleep spindles as regressors. The purpose of including IEDs and sleep spindles was to account for their potential influence on the hemodynamic response related to slow waves.

Beta-maps obtained at the group level after cluster-based correction for multiple comparison (p<0.05 cluster-wise significance threshold) are shown in Figure 3a (also see Table 2). Figure S1 and Table S1 represent uncorrected results. The occurrence of sleep slow waves was associated with a significant decrease in the hemodynamic response in two cortical clusters including left and right somatomotor regions (Table 2). The average signal temporal profiles for these regions showed that the maximum peak of the negative response was reached 4-8 seconds after slow-wave-onset (t=0) and was preceded by an increase in the hemodynamic signal that peaked ∼8 seconds before slow-wave onset (Figure 3b). No significant clusters were found at the subcortical level. No cortical or subcortical regions showed significant positive hemodynamic signal changes.

**Table 2.** Brain areas showing a significant BOLD-signal change time-locked to the occurrence of sleep slow waves (p< 0.05 after cluster correction). The table includes for each area the number of voxels and the coordinates of the center of mass in the standard MNI space.

| Signal Change | Region | Voxels | CM x | CM y | CM z |
| --- | --- | --- | --- | --- | --- |
| Negative | Right Somatomotor Cortex | 715 | -55.4 | +9.8 | +22.8 |
|  | Left Somatomotor Cortex | 487 | +49.5 | +16.7 | +42.8 |

**Figure 3.**
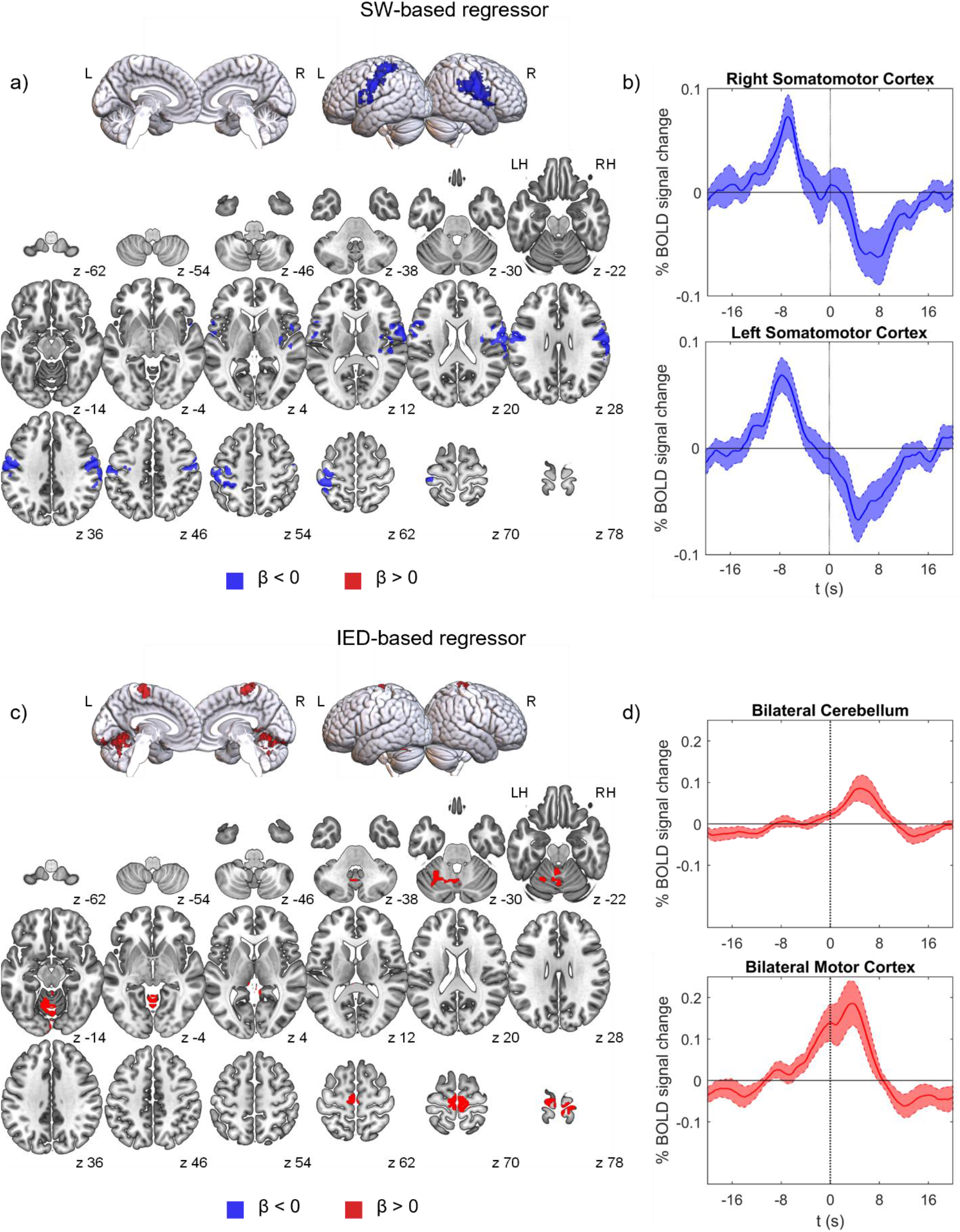
Results of the regression analysis after cluster-based correction for multiple comparison (p< 0.05, cluster forming threshold p<0.01, N=1000 permutations) for slow waves (a-b) and IEDs (c-d). Brain regions associated with a significant BOLD-signal increase are shown in red (β > 0), while regional BOLD-signal decreases are displayed in blue (β < 0). Brain images were generated using MRIcroGL (https://www.nitrc.org/projects/mricrogl/). Plots on the right show the mean BOLD-signals (up-sampled to the EEG sampling rate) time-locked to slow-waves (b) and to IEDs (d) onsets and the relative standard errors. These plots were obtained by averaging the signal of individual voxels within significant clusters identified through the regression analysis.

### Hemodynamic signal changes associated with spindles and IEDs

Spindles and IEDs were included in our model to account for their potential influence on the hemodynamic response related to slow waves. However, we also assessed their specific hemodynamic correlates.

We failed to detect any significant BOLD-signal changes associated with sleep spindles, either at the cortical or subcortical level. Results obtained using a liberal (uncorrected) statistical threshold are presented in Figure S2 and Table S2.

In line with previous observations obtained during wakefulness^36,61,62^, we found significant positive signal changes associated with the occurrence of IEDs in the bilateral motor cortex and bilateral cerebellum (Figure 3c-d; Table 3; Figure S3; Table S3). No cortical or subcortical regions showed negative hemodynamic changes.

**Table 3.** Brain areas showing a significant BOLD-signal change associated with IEDs density variations (p< 0.05 after cluster correction). The table includes for each area the number of voxels and the coordinates of the center of mass in the standard MNI space.

| Signal Change | Region | Voxels | CM x | CM y | CM z |
| --- | --- | --- | --- | --- | --- |
| Positive | Bilateral Cerebellum | 455 | +5.0 | +60.8 | -14.6 |
|  | Bilateral Motor Cortex | 257 | -0.9 | +24.8 | +72.9 |

### Comparison of cortical slow-wave-related hemodynamic changes in children and adults

Previous works using EEG demonstrated that the peak of slow-wave activity moves from posterior to anterior electrodes during development, mirroring the posterior-to-anterior gradient of cortical maturation^15,25,26,28^. To investigate the hemodynamic correlates of this EEG change, we generated a conjunction map and performed qualitative comparisons (Figure 4) between fMRI cortical activity maps (p<0.01, uncorrected) obtained in school-age-children and those obtained in a previous study on healthy adults^8^. We found that in both children and adults hemodynamic signal changes mainly occurred bilaterally in cortical areas around the central sulcus, with the largest overlap within the somatomotor cortex (Figure 4d). However, the comparison also showed that signal changes in children tended to extend more posteriorly, within the parietal cortex, while those of adults tended to extend more anteriorly, within frontal areas (Figure 4c).

**Figure 4.**
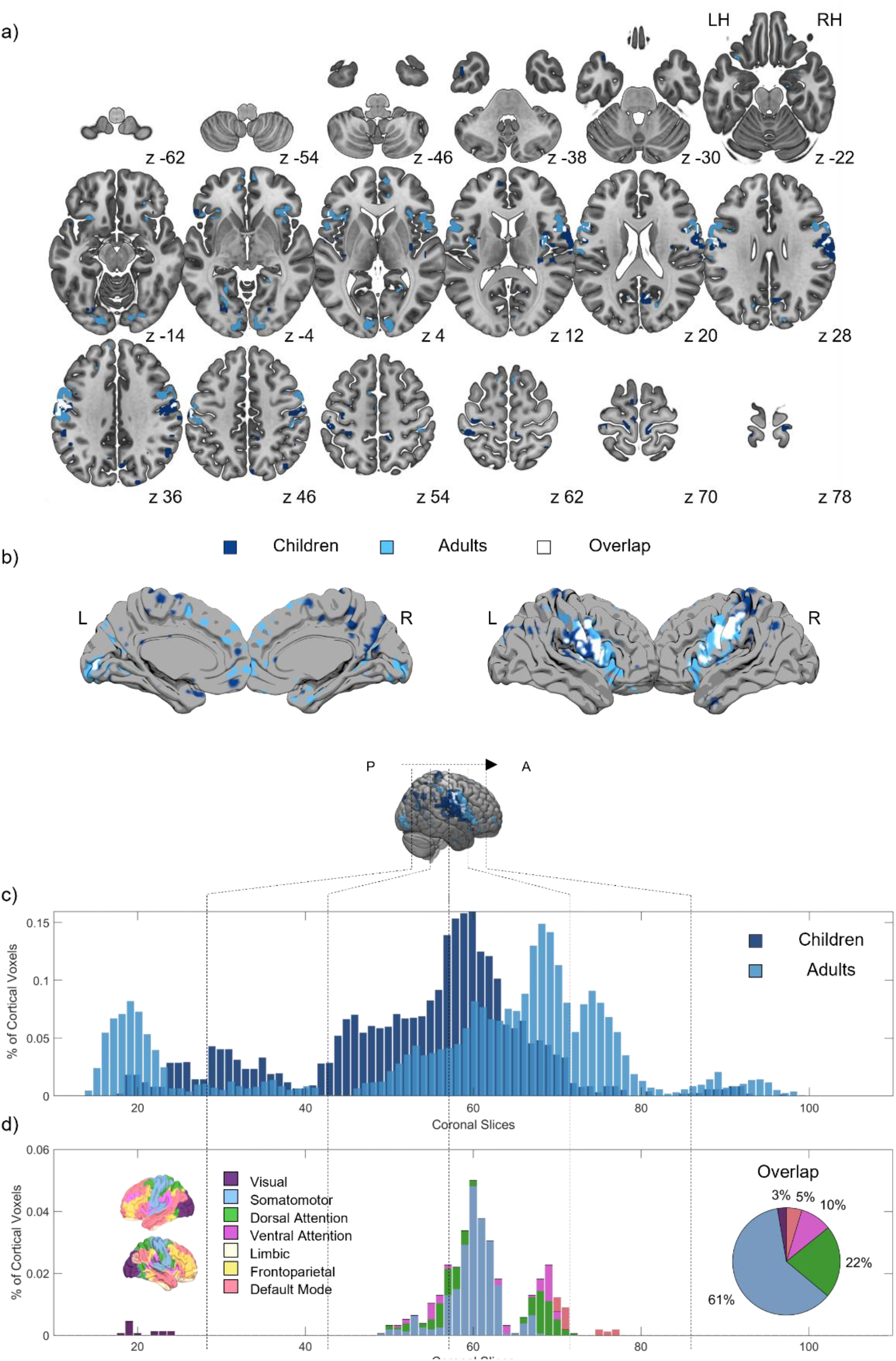
Conjunction maps showing hemodynamic changes associated with sleep slow waves in children (dark blue) and adults (light blue). Only cortical voxel with β < 0 and p < 0.01 are displayed for the two groups. Areas of overlap between children and adults are shown in white color. Panel a) shows multiple axial slices while panel b) presents brain surface plots generated using the Surf Ice software (https://www.nitrc.org/projects/surfice/). Panel c) shows the percentage of cortical voxels belonging to each consecutive coronal slice of the dataset, in the posterior to anterior direction separately for adults and children. Bars are plotted as semi-transparent. Panel d) shows the percentage of cortical voxels in each coronal slice for the area of overlap between children and adults. In this case, voxels belonging to distinct canonical functional networks are plotted separately and shown using a different color59,63. The pie plot reports the percentage of voxel from the overlap map belonging to each network. Most voxels appeared to be part of the somatomotor network (61%), followed by the dorsal and ventral attention networks.

### Effect of age on cortical regional involvement during sleep slow waves

Since slow waves travel at cortical level through cortico-cortical connections, they might involve multiple brain areas at different relative delays. Given that connectivity changes during development, the pattern of slow-wave cortical distribution may also be expected to change. We thus investigated the impact of age on the involvement of distinct cortical networks during sleep slow waves and found that while most networks did not exhibit any significant developmental changes, a positive correlation emerged between the coherence of BOLD profiles within DMN and age (R = 0.63, p<0.005, CI = 0.23-1.0, SE = 0.20; Figure 5). In other words, areas within the DMN showed an increasing degree of signal-change similarity during EEG slow waves as a function of age. This increased similarity in older children reflects the appearance of a more defined negative deflection in the average BOLD profile of the DMN (Figure S4 and Figure S5).

**Figure 5.**
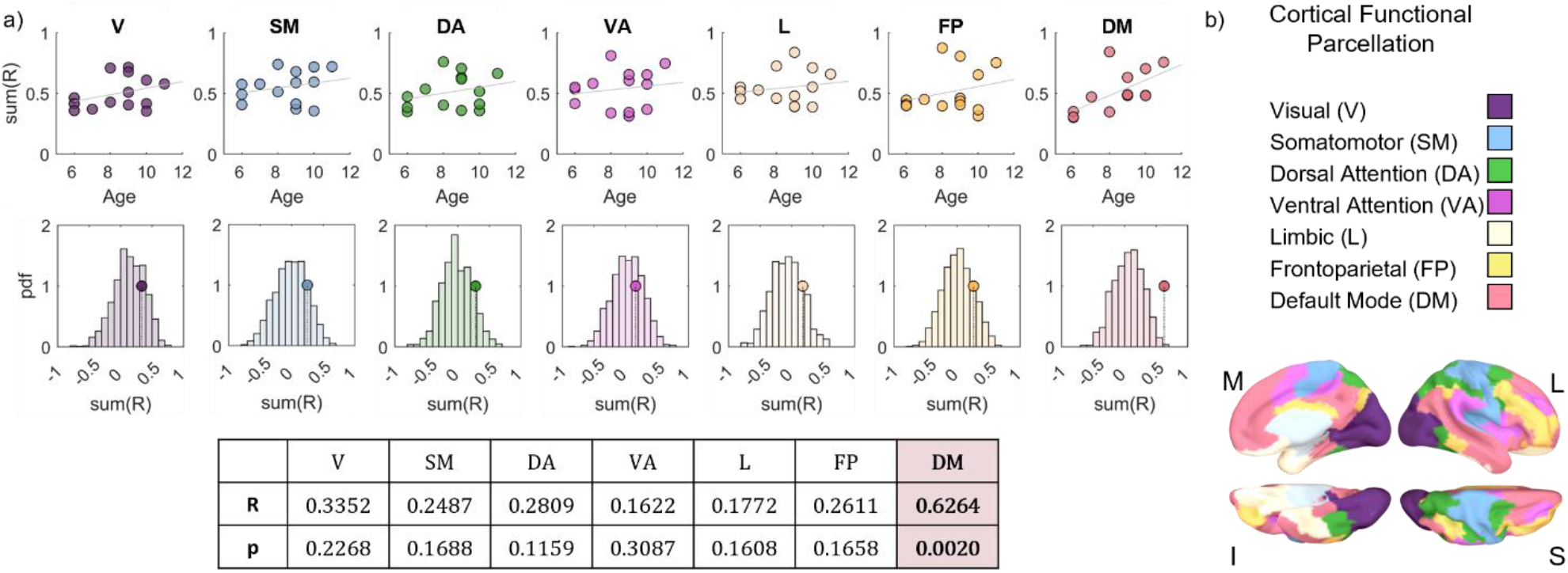
Correlation between cortical hemodynamic changes in the seven canonical functional networks and age. Panel a) shows for each cortical network the results of the correlation analysis between global similarity among BOLD profiles of regions belonging to the same network and age. Specifically, we first evaluated the mean (average across waves and voxels) BOLD profiles time-locked to sleep slow waves within 200 ROIs of the Schaefer atlas58. For each subject, we then estimated the correlation matrix among BOLD profiles of ROIs belonging to the same network and estimated a measure of similarity defined as the mean of the positive correlation coefficients derived from all the pairwise comparisons. Finally, the correlation between within-network similarity and age was calculated for each of the canonical networks (DA= Dorsal Attention; L= Limbic; DM= Default Mode; SM= Somato Motor; V= Visual; VA= Ventral Attention; FP= Fronto Parietal). For each network, the first row shows scatter plots including age and similarity score, while the second row displays the value of the correlation between similarity score and age relative to a null distribution of Pearson’s R-scores obtained from a permutation procedure (N= 1000). The DMN was the only network showing significant results after correction for multiple comparisons. Panel b) illustrates the cortical parcellation based on the seven canonical functional networks59.

### Correlation between thalamic slow-wave-related hemodynamic changes and age

While previous studies in adult individuals are consistent with an involvement of the thalamus in slow-wave expression, we found no group-level evidence of thalamic activity during slow waves in school-age children. This difference in thalamic response between children and adults could possibly reflect the occurrence of maturation-dependent modifications. The analysis, based on a subdivision of the thalamus into 7 ROIs according to their preferential connectivity to one of the canonical cortical networks^59^, revealed significant positive correlations of age with slow-wave-related hemodynamic changes for dorsal attention (R = 0.61, p = 0.002, CI = 0.33-0.90, SE = 0.15), limbic (R = 0.65, p = 0.002, CI = 0.27-1.02, SE = 0.19) and default mode networks (R = 0.68, p = 0.002, CI = 0.42-0.94, SE = 0.13; Figure 6). In line with these results, two adolescent subjects (age 15 and 17 y) with SeLECTS showed a pattern of thalamic response during slow waves that was more similar to that of adults than to that of younger school-age children. The results of a similar correlation analysis based on a thalamic (medial-anterior) ROI that showed significant positive BOLD-signal changes during slow waves in adults (for details see 8) are reported in Figure S6 (R = 0.71, p < 0.001, CI = 0.43-0.99, SE = 0.14).

**Figure 6.**
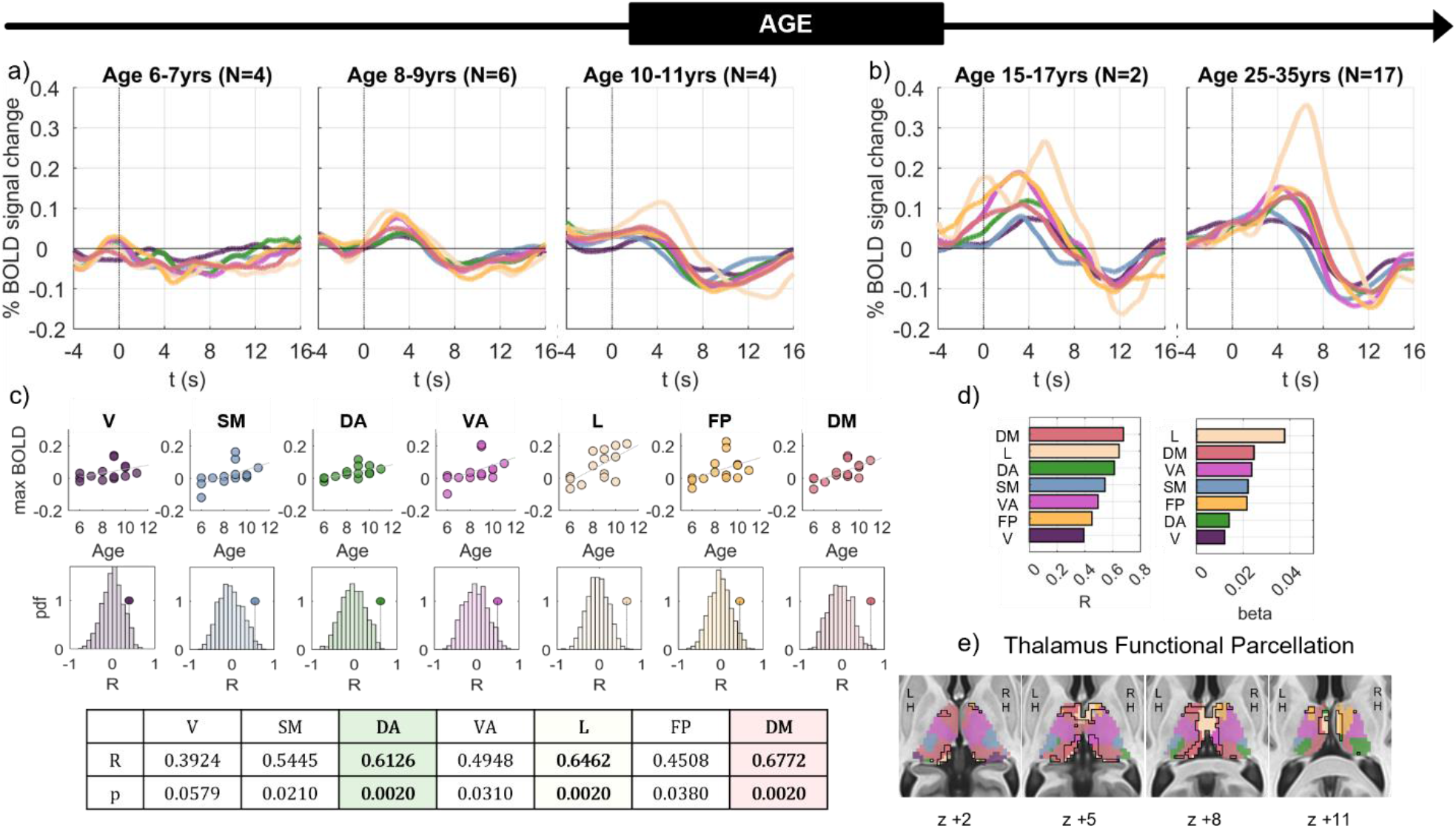
Correlation between thalamic hemodynamic changes and age. Panel a) shows the mean BOLD profiles time-locked to slow-wave onset for the seven thalamic ROIs in three sub-groups of children with increasing age, namely 6-7 years (4 subjects), 8-9 years (6 subjects), 10-11 years (4 subjects). Panel b) shows the mean BOLD profiles for 2 epileptic adolescents (15-17 years) and 17 healthy adults (25-35 years) included in our previous work^8^. Panel c) shows the results of the correlation between maximum amplitude of the BOLD-signal deflection and age with respect to the null-distribution obtained from the non-parametric permutation test for each of the seven thalamic ROIs. Only ROIs corresponding to the dorsal attention, limbic and default mode networks showed significant results (Pearson’s R-scores and p-values are reported in the table and significant effects are highlighted with colors). DA= Dorsal Attention; L= Limbic; DM= Default Mode; SM= Somato Motor; V= Visual; VA= Ventral Attention; FP= Fronto Parietal. Panel d) reports correlation coefficients (left) and the slope of the regression line (right) sorted in descending order for the seven ROIs. Panel e) illustrates the thalamic parcellation based on the preferential functional connectivity of each voxel with the seven canonical cortical networks^59^. The thalamic portion found to be activated in association with the occurrence of slow waves in adults is highlighted and enclosed with a black line^8^.

## Discussion

The slow waves of NREM sleep are known to undergo relevant topographical changes from childhood to adulthood, paralleling brain maturational adaptations and the acquisition of cognitive and behavioral skills. The present work used simultaneous EEG-fMRI to investigate the cortical and subcortical hemodynamic correlates of sleep slow waves in school-age children and identify potential differences and similarities with respect to those of adults^8,9^. We found that the occurrence of slow waves is associated in both children and adults with significant negative hemodynamic changes in the somatomotor cortex, which however extend more posteriorly in children and more anteriorly in adults. Moreover, our results suggest that while slow waves involve on average a large portion of the cortex in both children and adults, the involvement of areas belonging to the DMN increases across development. Finally, while we did not observe significant hemodynamic changes during slow waves in subcortical structures, thalamic activity showed a positive correlation with age. Overall, these findings indicate that the somatomotor cortex has a key role in the expression of sleep slow waves across the lifespan and also suggest that the thalamic contribution to slow-wave regulation may change during development.

### The somatomotor cortex is involved in slow-wave expression in both children and adults

We found that slow waves of school-age children are associated with negative hemodynamic changes in the left and right somatomotor cortices. These regions showed substantial overlap with those we previously found in healthy adult individuals^8,9^. Interestingly, previous studies conducted on adults also showed that slow waves behave as traveling waves with a specific origin and propagation pattern and that the somatomotor area represents one of the main slow-wave origin hotspots^8,21–23^. Overall, these results indicate that the somatomotor cortex may have a special role in slow-wave generation and expression across the lifespan. But why is that so? Evidence indicates that slow waves originate more often from, and SWA is higher in, areas that are more intensively active during wakefulness prior to sleep^13,14^. Indeed, previous studies showed that experience-dependent plasticity induced through extended practice of motor, tactile, or visual tasks results in a local SWA increase within task-related areas13,64,65, whereas reduced experience-dependent activation (e.g., through arm immobilization or visual deprivation) causes opposite changes^14,66^. A possibility is thus that the somatomotor cortex may naturally accumulate more experience-dependent plastic adaptations during wakefulness and thus show a stronger tendency to generate slow waves in sleep. Other potential explanations might be suggested based on the relationship between the somatomotor cortex and arousal-related structures. Indeed, the somatomotor cortex contains the highest degree of noradrenergic innervation in the human and monkey cortex^67,68^ and relatedly shows strong BOLD-signal changes in correspondence with momentary variations in the level of arousal^69,70^. In this light, neurons of the somatomotor cortex could be especially vulnerable to changes in arousal and may have a stronger propensity to generate slow waves when activatory neurotransmitters are tonically reduced. Moreover, it has been previously suggested that transient activations of arousal-related structures targeting the somatomotor cortex may synchronize large neuronal populations favoring the emergence of large-amplitude slow waves, especially in this area^43,46^.

### BOLD-signal changes during slow waves partially differ between children and adults

Although the somatomotor cortex showed similar hemodynamic changes in children and adults^8^, patterns of changes showed qualitative differences between the two age groups. Indeed, in children, activity related to slow waves tended to involve more posterior cortical regions, and in particular parietal areas, while in adults they extended more anteriorly, involving frontal regions. This observation is consistent with previous findings based on the analysis of scalp EEG SWA, which showed a posterior-to-anterior shift from childhood to adulthood^2,24,33,71^. Such variations appear to mirror the posterior-to-anterior gradient of cortical brain maturation and the parallel acquisition of specific cognitive and behavioral skills^26,31,72^. Indeed, changes in regional slow-wave expression have been suggested to reflect in particular the consequences of synaptic pruning and refinement^24,73^.

### Involvement of DMN during slow waves changes with age

Previous work demonstrated that the propagation of slow waves at cortical level depends on the integrity of cortico-cortical connections^23^. During development, the distance traveled by slow waves appears to increase as a function of age^33,34,74^, reflecting the process of fiber myelination and the maturation of white-matter connectivity^33,72,75^. In fact, while local, short-range connections appear to predominate in children, brain maturation leads to the establishment of a more distributed, long-range connectivity pattern^76^. Moreover, the maturation of functional brain circuits appears to follow the hierarchical organization of the human brain, with the somatomotor network being the first to reach an adult-like architecture and the DMN being the latest among cortical networks^77–79^. In line with these considerations, our results demonstrated that activity of regions belonging to the DMN during sleep slow waves changes across development, becoming more coherent and consistent as a function of age. Such changes may depend on, and thus mirror, the maturation of functional and anatomical connectivity of the DMN^80–82^. Taken together, these findings support the use of sleep slow waves as a valuable, noninvasive marker able to track the state and modifications of regional and long-range connectivity in humans^33^.

### Thalamic activity associated with slow waves changes across development

While the sleep slow wave is regarded as a cortical phenomenon, studies on animal models revealed that its expression is at least partially influenced by subcortical structures, including the thalamus ^3– 5,83,84.^ In particular, the spontaneous firing of higher-order and centromedial thalamic nuclei is responsible for initiating and synchronizing the transition between off- and on-periods in cortical neuronal populations60,85,86. In addition, recent work performed on epilepsy patients who underwent simultaneous scalp EEG and deep thalamic recordings showed that slow waves in the anterior thalamic nuclei precede those detected at cortical level and suggested that the thalamus may thus have a direct role in orchestrating these cortical oscillations^7^. Further evidence supporting the involvement of the thalamus in slow wave activity was provided by studies using thalamic optogenetic stimulation, which resulted in the induction of cortical slow waves^4,5^. Consistent with these observations, non-invasive investigations through combined EEG-fMRI revealed changes in hemodynamic thalamic activity time-locked to sleep slow waves in healthy adults^8,9,87^. Surprisingly, though, we did not find a similar change in hemodynamic thalamic activity during sleep slow waves in school-age children with SeLECTS. However, we found a positive, significant correlation between the amplitude of thalamic hemodynamic changes and the age of children, and qualitatively showed that signal changes in two older adolescents were characterized by a response pattern that appeared to be intermediate between those of children and adults. Potentially, the observed changes in thalamic activity might reflect changes in the strength and organization of thalamo-cortical interactions during development^88–91^. Interestingly, the largest age-dependent changes were found in portions of the thalamus interacting with the cortical default mode, limbic, and dorsal attention networks. Of note, limbic and default mode portions of the thalamus were previously found to be among those showing the strongest hemodynamic modulation during slow waves in healthy adults^9^. The limbic thalamus is connected to important structures related to memory, such as the prefrontal cortex and hippocampus^92^. Moreover, hippocampal specific oscillations-the sharp-wave ripples (150-250 Hz)-along with thalamic sleep spindles and cortical slow waves, appear to support memory consolidation^11,93^. Thus, the observed changes in limbic-thalamus activity during slow waves might reflect, or even contribute to, the maturation of sleep-related memory consolidation processes^94^. On the other hand, changes in the activity of portions of the thalamus connected with the DMN could be (causally) related to the parallel changes observed in the involvement of this cortical network during sleep slow waves^95^.

Overall, these observations indicate that changes in thalamic activity in association with slow-wave occurrence, and thus, potentially, the involvement of the thalamus in regulating slow-wave expression, change significantly from childhood to adulthood.

### Limitations

Present analyses were performed on EEG-fMRI data collected from children with self-limited focal epilepsy who fell asleep during afternoon resting-state scan sessions. The epileptic syndrome and related antiepileptic medications may have determined alterations in sleep (micro)structure and/or hemodynamic changes associated with slow waves, especially in perisylvian areas where IEDs mainly occur^96^. While we carefully assessed possible interactions between IEDs and slow waves and found no evidence of significant confounding effects, future studies in children without concomitant pathological conditions will be necessary to validate the present findings. Moreover, since data collection was performed during the afternoon, we cannot exclude possible circadian or homeostatic differences relative to typical overnight sleep. However, previous studies investigating the hemodynamic correlates of slow waves in adults obtained very similar results using data collected overnight^9^ or during an afternoon nap^8^.

While previous work showed that sleep spindles are associated with BOLD-signal changes in the thalamus^97–103^, here no significant effects were observed at both the cortical and subcortical levels. This may be due to insufficient statistical power, as relatively few spindle events were entered into the regression analysis, but it could also reflect an alteration of thalamic activity related to the underlying pathological condition of our patients.

Analyses based on cortical ROIs used parcellations obtained from functional connectivity estimates in adult individuals^8,59^. Since cortico-cortical and thalamo-cortical connectivity is known to partially differ between children and adults due to maturational adaptations^104^, our choice might have negatively affected the accuracy of labels assigned to cortical and thalamic sub-regions in the sample of school-age children. On the other hand, this approach allowed us to more directly compare results obtained in children with those previously described in adult individuals using similar methods and analyses^8^.

Notwithstanding the above limitations, it should be kept in mind that studies on underage populations pose relevant ethical problems, especially if involving experimental protocols combining different techniques, as in the present work. In this light, while the discussed limitations warrant caution regarding generalizability to healthy children and physiological, overnight sleep, we hope that this study will represent the basis for future investigations aimed at determining developmental changes in the contribution of cortical and subcortical structures to the establishment of physiological sleep rhythms and (micro)structure.

## Acknowledgements

This work was supported by intramural funds from the IMT School for Advanced Studies Lucca (to G.B., M.B., E.R., G.H., D.B), and a grant “Dipartimenti di Eccellenza 2018–2022″, MIUR-Italy, to the Department of Biomedical, Metabolic and Neural Sciences (S.M. and F.B.).

## Conclusions

Present findings revealed that the cortical and subcortical correlates of sleep slow waves present both similarities and differences with respect to those observed in healthy adult individuals. At the cortical level, our results are in line with previous findings indicating that the distribution of SWA changes during development, shifting from posterior to anterior brain areas^25^. Moreover, we demonstrated that involvement of regions belonging to the DMN in sleep slow waves increases during development, likely mirroring the structural and functional maturation of this particular network. On the other hand, our results also indicated that the somatomotor cortex may represent a key hub for the generation and expression of sleep slow waves in both children and adults, and thus, potentially, throughout the entire lifespan^8,9,87^. Finally, we found that thalamic involvement related to the occurrence of slow waves changes during development, being absent in early childhood and increasing progressively with age. This observation suggests that developmental changes in slow-wave characteristics may not depend only on the maturation of cortical gray matter and cortico-cortical connections. Indeed, the maturation of the thalamus and thalamo-cortical connectivity might also have a crucial role in determining physiological maturation-dependent changes in the regulation and expression of NREM slow waves. Advancing our knowledge of such changes could have fundamental implications for understanding how the cortex and subcortical structures interact to generate sleep rhythms and their essential functions.

