## Supplemental figures and tables for "Maturation-dependent changes in cortical and thalamic activity during slow waves of light sleep: insights from a combined EEG-fMRI study"

***Figure S1***

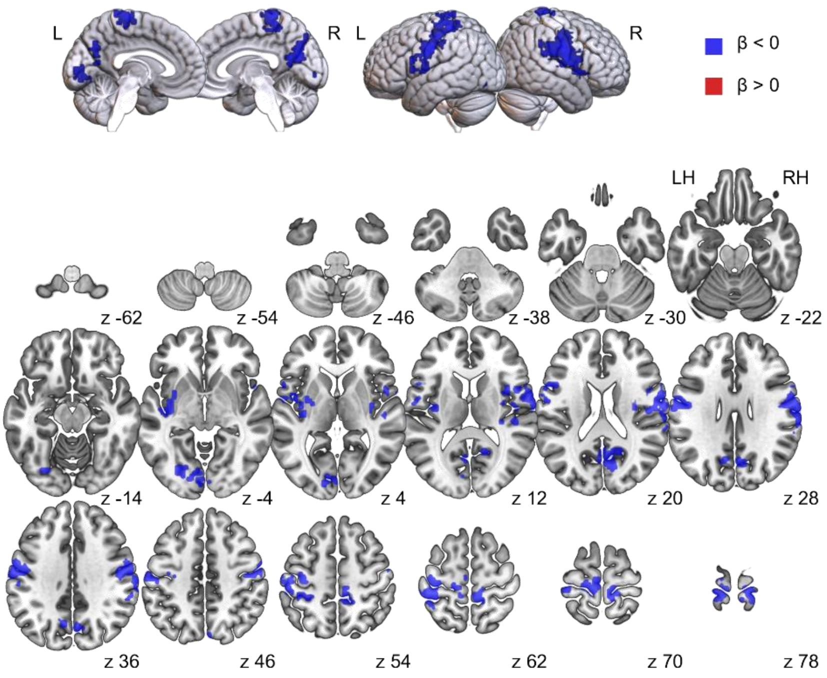


*Figure S1. Results of the regression analysis before cluster correction (p <0.01, cluster size 50 voxels) for slow waves. BOLD-signal increases (>0) are shown in red color while BOLD signal decreases (<0) are represented in blue. Brain images were generated using MRIcroGL ([https://www.nitrc.org/projects/mricrogl](https://www.nitrc.org/projects/mricrogl)).*

***Figure S2***

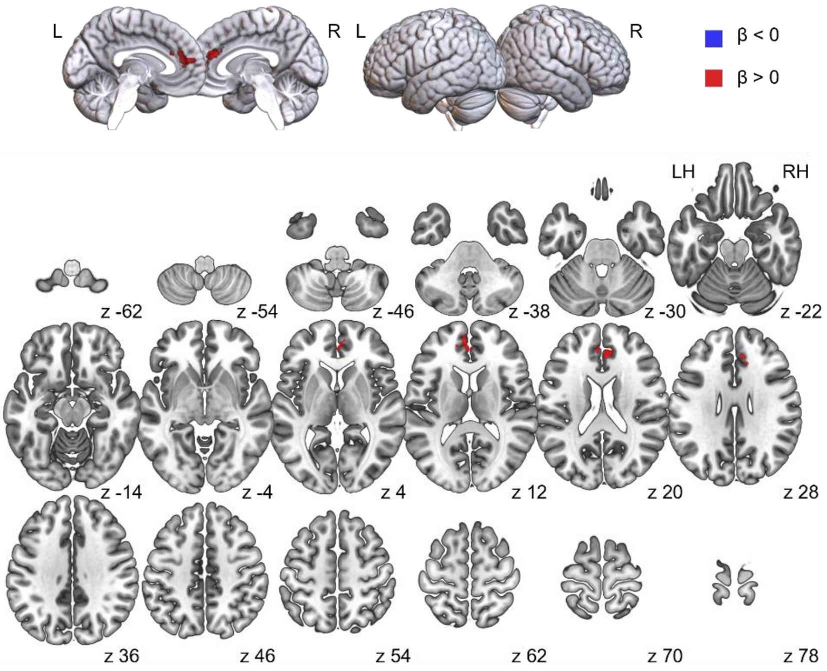


*Figure S2. Results of the regression analysis before cluster correction (p <0.01, cluster size 50 voxels) for spindles. BOLD-signal increases (>0) are shown in red color while BOLD signal decreases (<0) are represented in blue. Brain images were generated using MRIcroGL (https://www.nitrc.org/projects/mricrogl).*

***Figure S3***

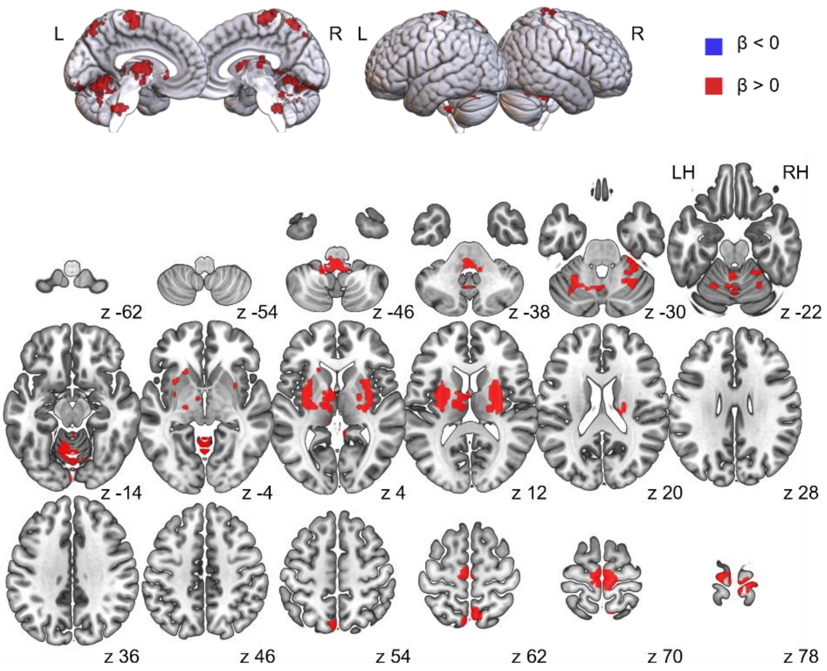


*Figure S3. Results of the regression analysis before cluster correction (p <0.01, cluster size 50 voxels) for IEDs. BOLD-signal increases (>0) are shown in red color while BOLD signal decreases (<0) are represented in blue. Brain images were generated using MRIcroGL (https://www.nitrc.org/projects/mricrogl).*

### *Table S1*

| Signal Change | Region | Voxels | CM x | CM y | CM z |
|---|---|---|---|---|---|
| Negative | Right Somatomotor Cortex | 744 | -55.0 | +9.6 | +22.6 |
| | Left Somatomotor Cortex | 487 | +49.6 | +16.4 | +42.4 |
| | Precuneus | 192 | -4.3 | +66.1 | +25.6 |
| | Right Paracentral | 123 | -12.6 | +35.3 | +68.7 |
| | Left Paracentral | 108 | +13.2 | +29.6 | +71.7 |
| | Left Insular Cortex | 78 | +37.7 | +9.7 | +3.5 |
| | Left Visual Cortex | 76 | +14.2 | +82.8 - | -2.3 |

*Table S1. Brain areas showing a significant BOLD-signal change time-locked to the occurrence of sleep slow waves before cluster correction (p <0.01, cluster size 50 voxels). The table includes for each area the number of voxels and the coordinates of the center of mass in the standard MNI space.*

### *Table S2*

| Signal Change | Region | Voxels | CM x | CM y | CM z |
|---|---|---|---|---|---|
| Positive | Anterior Cingulate Cortex | 120 | -0.9 | -42.6 | +17.4 |

*Table S2. Brain areas showing a significant BOLD-signal change time-locked to the occurrence of sleep spindles before cluster correction (p <0.01, cluster size 50 voxels). The table includes the number of voxels for each area and the coordinates of the center of mass in the standard MNI space.*

***Table S3***

| Signal Change | Region | Voxels | CM x | CM y | CM z |
|---|---|---|---|---|---|
| Positive | Left Cerebellum | 457 | +4.4 | +60.7 | -14.7 |
| | Bilateral Motor Cortex | 265 | -0.7 | +24.2 | +72.7 |
| | Left Thalamus and Basal Ganglia | 245 | +22.5 | +5.9 | +6.7 |
| | Right Basal Ganglia | 177 | -27.5 | +6.7 | +9.1 |
| | Bilateral Parietal Cortex | 98 | -0.4 | +65.0 | +62.8 |
| | Brainstem (Medulla) | 66 | +3.0 | +39.2 | -42.3 |
| | Right Cerebellum | 59 | -28.1 | +46.0 | -27.1 |

*Table S3. Brain areas showing a significant BOLD-signal change associated with IED density variation before cluster correction (p <0.01, cluster size 50 voxels). The table includes the number of voxels for each area and the coordinates of the center of mass in the standard MNI space.*

***Figure S4***

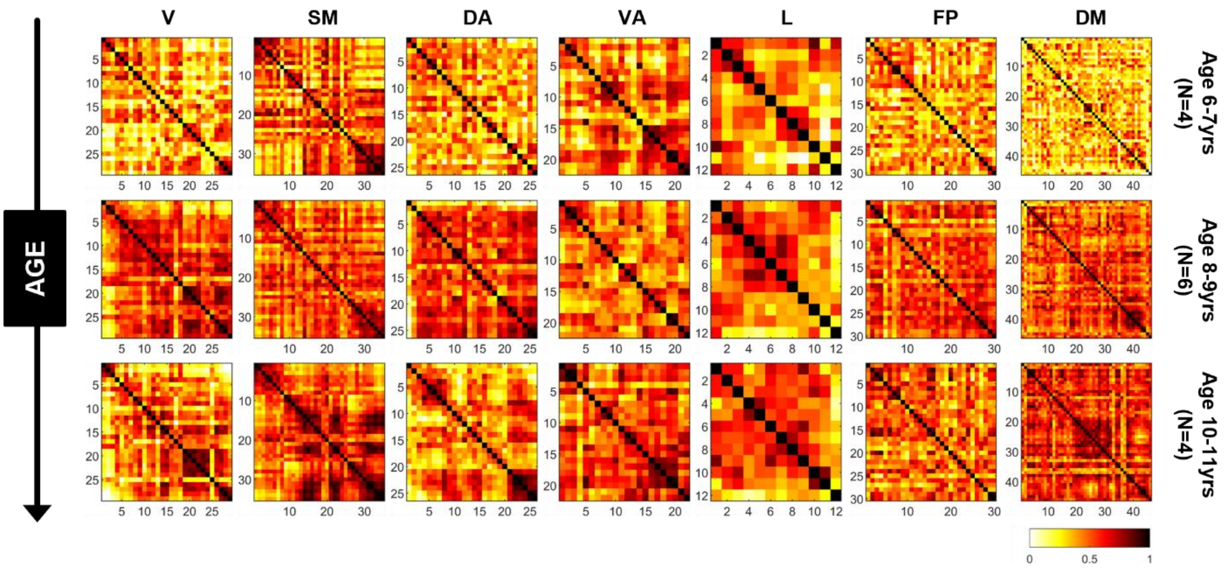


*Figure S4. Changes in within-network correlation matrices as a function of age. We first computed the mean (average across waves and voxels) BOLD profiles time-locked to sleep slow waves within 200 ROIs of the Schaefer atlas. Then, for each subject, we estimated the correlation matrix among BOLD profiles of ROIs belonging to the same canonical cortical networks (DA= Dorsal Attention; L= Limbic; DM= Default Mode; SM= Somato Motor; V= Visual; VA= Ventral Attention; FP= Fronto Parietal). In order to qualitatively illustrate how activity changed as a function of age, average correlation matrices are displayed for three age sub-groups: 6-7 years (N=4), 8-9 years (N=6), 10-11 years (N=4).*

### *Figure S5*

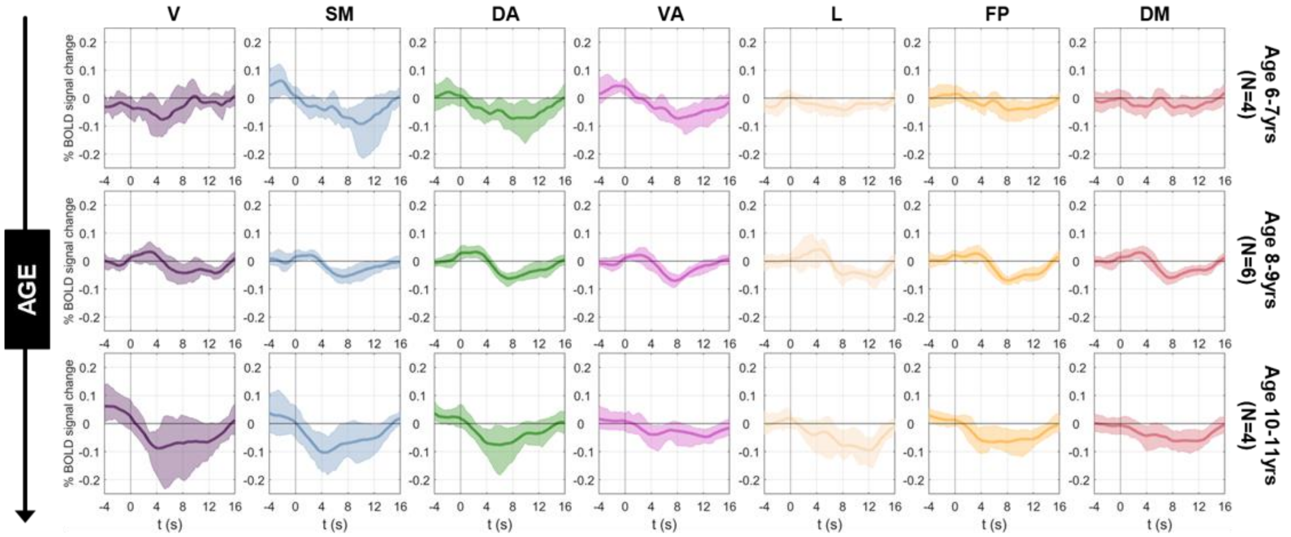


*Figure S5. Changes in cortical BOLD profiles time-locked to sleep slow waves as a function of age. We first computed the mean (average across waves, voxels and subjects) BOLD profile within 200 spatial ROIs of the Schaefer atlas, and then further averaged the signals of ROIs belonging to the same canonical cortical networks: DA= Dorsal Attention; L= Limbic; DM= Default Mode; SM= Somato Motor; V= Visual; VA= Ventral Attention; FP= Fronto Parietal. Plots are presented for three age sub-groups: 6-7 years (N=4), 8-9 years (N=6), 10-11 years (N=4). In order to qualitatively illustrate how activity changed as a function of age, average correlation matrices are displayed for three age sub-groups: 6-7 years (N=4), 8-9 years (N=6), 10-11 years (N=4). Colored shadows represent for each time-point the range containing the 75% of ROI profiles around the mean.*

### *Figure S6*

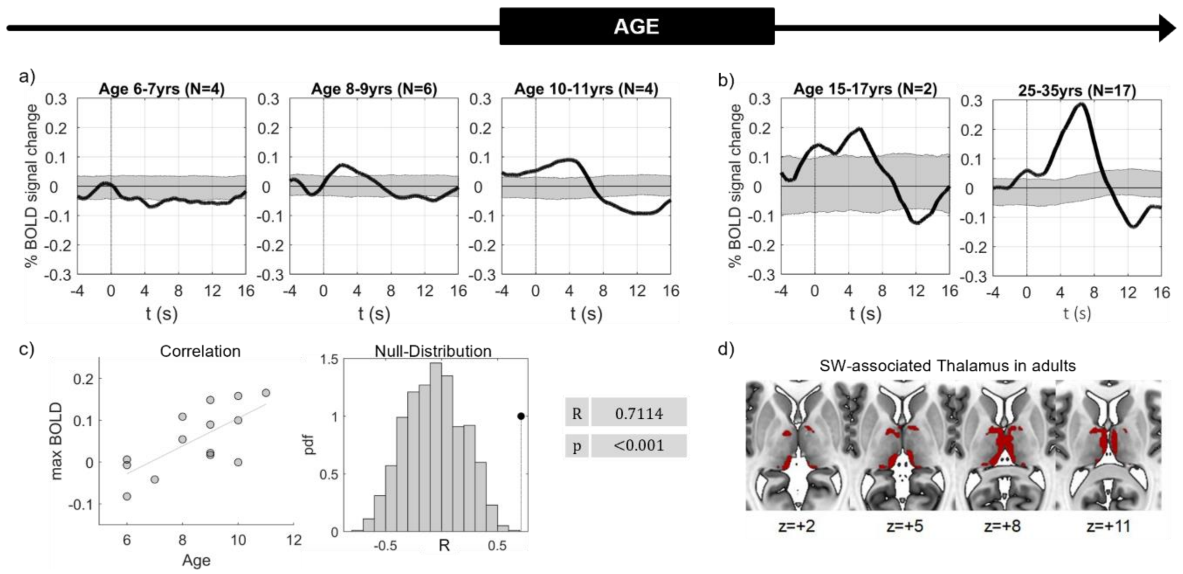


*Figure S6. Correlation between thalamic hemodynamic changes during sleep slow waves and age. This analysis was focused on the portion of the thalamus that was previously found to be activated in association with the occurrence of sleep slow waves in a group of healthy adults. The upper row of panel a) shows the mean BOLD profiles time-locked to slow wave onset (t=0) in three sub-groups of children with increasing age, namely 6-7 years (4 subjects), 8-9 years (6 subjects), 10-11 years (4 subjects) group. The plots show that the positive BOLD change following slow-wave occurrence becomes more pronounced with increasing age. The gray shadow reports the 5th-95th percentile range of the 1000 correspondent mean BOLD profiles obtained from the permutation procedure (N=1000). Panel b) reports in a similar manner the mean BOLD profiles for a group of two 2 epileptic adolescents (15-17 years) and a group of 17 25-35 years old healthy adults (25-35 years)[8]. Panel c) shows BOLD maximum amplitude values as a function of subject age and the results of the correlation analysis compared to the null-distribution obtained from the non-parametric permutation test,*

*where we shuffled the time onset of slow waves across subjects (N=1000). After correction, the p-value remained below the standard significance threshold ($p<0.01$). The results revealed a positive correlation between age and the maximum percentual BOLD average signal, even accounting for the density of the IEDs. Panel d) illustrates topographically on axial brain slices the thalamic portion considered in the analysis*[8].
